# Dynamic properties of internal noise probed by modulating binocular rivalry

**DOI:** 10.1101/480988

**Authors:** Daniel H. Baker, Bruno Richard

**Author notes:** Corresponding author*: Daniel H Baker, Department of Psychology, University of York, Heslington, York, United Kingdom, YO10 5DD.

## Abstract

Neural systems are inherently noisy, and this noise can affect our perception from moment to moment. This is particularly apparent in binocular rivalry, where our perception of competing stimuli shown to the left and right eyes alternates over time in a seemingly random fashion. We investigated internal noise using binocular rivalry by modulating rivalling stimuli using dynamic sequences of external noise of various rates and amplitudes. As well as measuring the effect on dominance durations, we repeated each external noise sequence twice, and assessed the consistency of percepts across repetitions. External noise modulations with standard deviations above 4% contrast increased consistency scores above baseline, and were most effective at 1/8Hz. A computational model of rivalry in which internal noise has a 1/f (pink) temporal amplitude spectrum, and a standard deviation of 16%, provided the best account of our data, and was able to correctly predict perception in additional conditions. Our novel technique provides detailed estimates of the dynamic properties of internal noise during binocular rivalry, and by extension the stochastic processes that drive our perception and other types of spontaneous brain activity.

**Significance statement:** Although our perception of the world appears constant, sensory representations are variable because of the ‘noisy’ nature of biological neurons. Here we used a binocular rivalry paradigm, in which conflicting images are shown to the two eyes, to probe the properties of this internal variability. Using a novel paradigm in which the contrasts of rivalling stimuli are modulated by two independent external noise streams, we infer the amplitude and character of this internal noise. The temporal amplitude spectrum of the noise has a 1/f spectrum, similar to that of natural visual input, and consistent with the idea that the visual system evolved to match its environment.

## Introduction

Despite appearing constant, our sensory perception fluctuates from moment to moment because of the non-deterministic nature of biological neurons. This ‘internal noise’ operates at multiple timescales, and affects our decisions about sensory information. Internal noise is particularly apparent in bistable phenomena such as binocular rivalry, in which our perception of conflicting images shown to the two eyes fluctuates over time in a stochastic fashion. Because phenomena like rivalry make otherwise invisible processes available to conscious perception, they provide a useful tool for probing the properties of internal noise.

In a typical rivalry experiment, participants view sine wave grating patterns with orthogonal orientations in the left and right eyes (see Figure 1a). They are asked to report which stimulus they perceive at each moment in time by continuously pressing a response button that corresponds to the perceived orientation. Histograms of the durations for which each image remains dominant typically have positive skew, approximating a gamma distribution (or a normal distribution on logarithmic axes). Computational models of rivalry (e.g. Kim et al., 2006; Lehky, 1988; Wilson, 2007, 2003) have successfully explained the statistical pattern of percepts reported by assuming the presence of three key processes: inhibition between neurons representing the two stimuli, adaptation to the dominant stimulus, and noise. Inhibitory properties have been investigated using dichoptic masking paradigms (Baker and Meese, 2007; Legge, 1979; Meese and Baker, 2009) and by varying the properties of rivalling stimuli (Baker and Graf, 2009a, 2009b; Stuit et al., 2011, 2009), and there is direct evidence of adaptation during a period of dominance (Alais et al., 2010). However, comparatively little is known about the precise properties of the noise, as there have been few attempts to investigate it directly, despite recognition of its importance (Brascamp et al., 2006; Lehky, 1995; Moreno-Bote et al., 2007; Shpiro et al., 2009).

**Figure 1:**
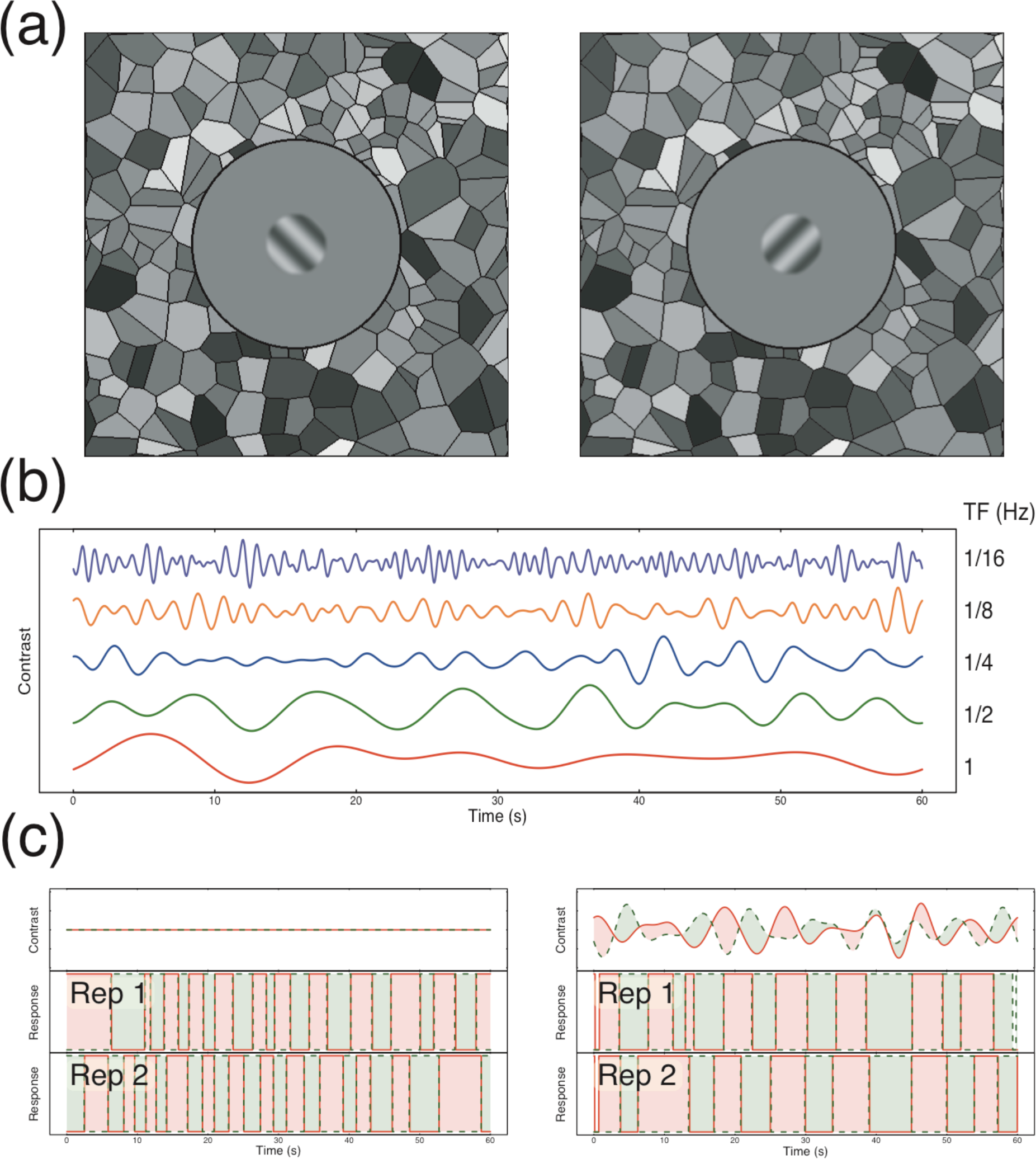
Methodological details. Panel (a) shows example stimuli with conflicting orientations, surrounded by a Voronoi texture to aid binocular fusion. Panel (b) shows example waveforms used to modulate stimulus contrasts at the five temporal frequencies used in the experiment. Panel (c) shows example trial timecourses for two repetitions of an unmodulated condition (left) and a modulated condition (right). Red (green) regions in the lower two plots indicate periods of time when the left (right) eye’s stimulus was perceived. Note that in the example on the right, percepts closely followed the physical contrast with a slight lag.

One exception is a study that randomly manipulated the coherence of rivalling dot motion stimuli throughout a trial in order to influence alternations (Lankheet, 2006). By reverse-correlating coherence with the observers’ percepts, a biphasic profile was apparent, in which coherence was stronger in the suppressed eye and weaker in the dominant eye during the ~1s preceding a flip. This pattern was reversed at longer pre-flip durations, and overall the results were predicted by a simple rivalry model featuring adaptation and mutual inhibition. Although the results demonstrate that external noise can influence rivalry alternations, the parameters of the external noise were not manipulated, and so the results can reveal little about the characteristics of internal noise.

Other work has aimed to influence rivalry alternations by periodically changing the contrast of the rivalling stimuli. In a study by O’Shea and Crassini (1984), the contrasts of rivalling gratings were periodically reduced to 0, either in phase or in antiphase across the eyes. At modulation frequencies above 20Hz (and sometimes as low as 3Hz), rivalry alternations still occurred as normal regardless of phase, suggesting a persistance in the underlying mechanism (see also Buckthought et al., 2008; Leopold et al., 2002). In a related study, Kim, Grabowecky and Suzuki (2006) used a square wave temporal modulation to alter the contrast of rivalling stimuli in antiphase (i.e. one stimulus increased in contrast and the other decreased at the same time) at a range of temporal frequencies from 0.28Hz to 2.48Hz. This manipulation caused a peak in the histogram of dominance durations at the half period of the modulation frequency. The increase was greatest when the half period was 600ms, a duration corresponding to the peak of the histogram for unmodulated rivalry using the same stimuli. Furthermore, there were additional peaks at odd integer harmonics of the modulation frequency. The authors consider this to be evidence of a stochastic resonance effect, and support this with a computational model of rivalry alternations.

Here we extend these approaches by modulating the contrast of rivalling stimuli using two independent dynamic noise sequences instead of square wave modulations (see Figure 1b,c). As well as measuring the effect on dominance durations, this design allows us to reverse correlate the participant’s reported percept with the timecourse of the external noise. In addition, we can use the same noise sequences multiple times, and measure the consistency of the participants’ percepts in a dynamic version of the ‘double pass’ paradigm (Burgess and Colborne, 1988; Green, 1964). If the external noise sequences were entirely determining perception, responses should be identical across the two repetitions. On the other hand, if the external noise sequences have no influence on perception then the similarity of responses will be determined by internal noise, and response consistency will be that expected by chance. The empirically measured consistency scores will therefore give an index of the relative influences of internal and external noise on perception. By manipulating the variance and temporal frequency content of the noise sequences, we can investigate properties of the internal noise that influences rivalry alternations. We interpret the results with reference to an established computational model of rivalry proposed by Wilson (2007, 2003) (see Figure 2), to which we add different types of internal noise.

**Figure 2.**
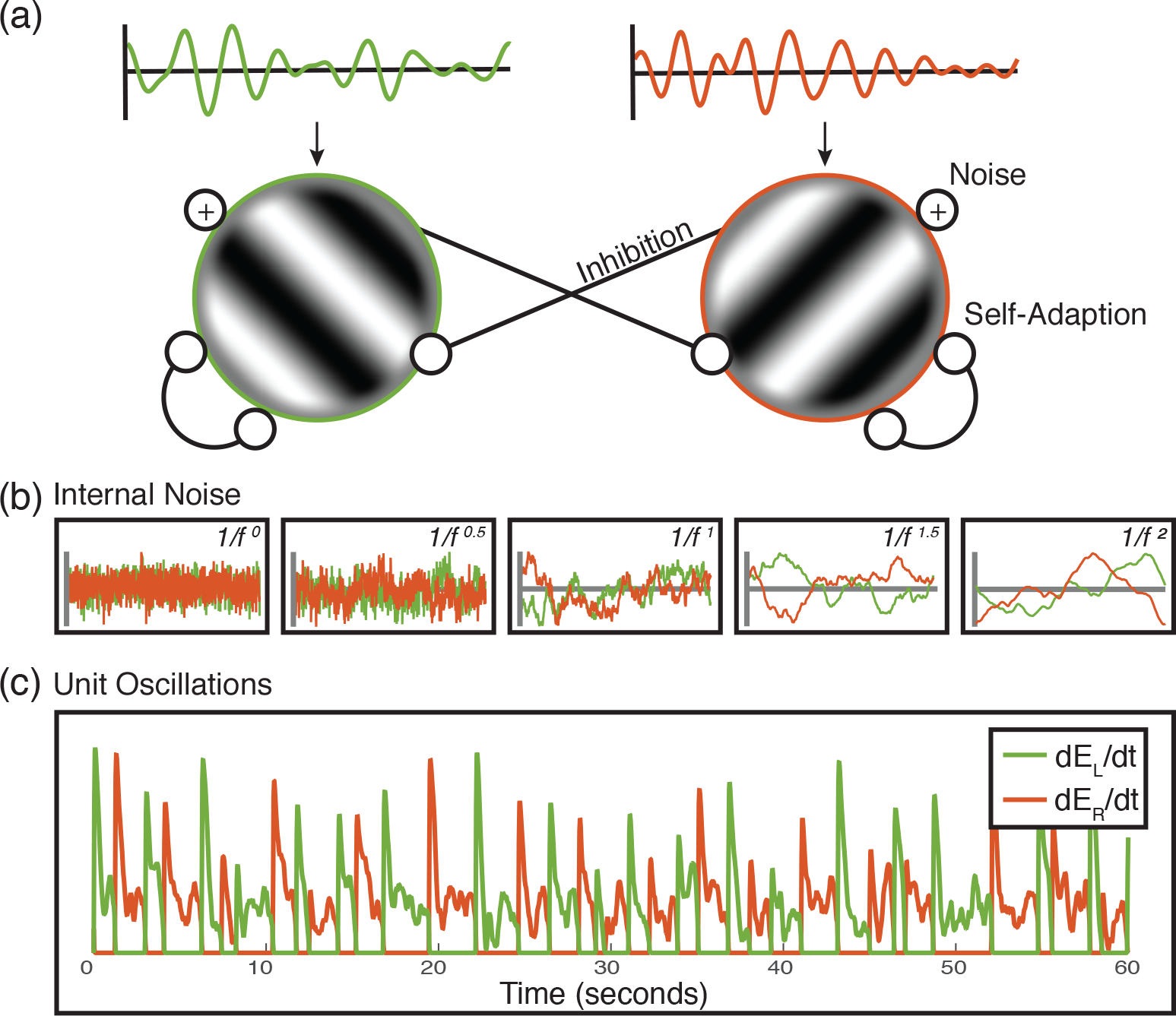
Model details. (a) Model diagram of the two competing units. Each receives as input an independent white noise stream, bandpass filtered at one of five different temporal frequencies (see Methods). The minimum rivalry model (Wilson, 2007) defines the oscillatory behaviour of rivalry between two units with self-adaptation and mutual inhibition. We include additive internal independent monocular noise in our model, marked by the (+) symbol. (b) Examples of the five different internal noise spectral slopes (α = 0 − 2.0) of the model for the left (green) and right (red) responding units. Noise streams with steeper slopes have an increased relative amplitude of low temporal frequencies relative to high, which leads to slower changes in the noise amplitude. (c) Example oscillatory behaviour of the model for a given trial (60s). The colours represent the responses of the left (green) and right (red) responding units.

## Results

### External noise strongly modulates binocular rivalry alternations

In the absence of any noise modulations, binocular rivalry produced a typical histogram of dominance durations with a positive skew (see grey curve in Figure 3ai), and a mean of 2.7 seconds. A 5 × 5 repeated measures ANOVA indicated that the mean dominance duration depended on both temporal frequency (F(4,16)=34.43, *p*<0.001, *η*_*p*_^2^=0.90) and modulation contrast (F(4,16)=8.15, *p*<01, *η*_*p*_^2^=0.67), and also showed that the two variables interacted (F(16,64)=18.01, *p*<0.001, *η*_*p*_^2^=0.82). The histograms in Figure 3a show that at lower temporal frequencies, strong contrast modulation resulted in slightly more long-duration percepts (an increase in positive skew), whereas at higher temporal frequencies the peak of the histogram shifted leftwards. These patterns were reflected in both the change in means (Figure 3b) and also the shift in the autocorrelation functions (Figure 3c), such that high temporal frequencies (e.g. the purple curve) had a shorter lag than long ones (e.g. the red curve). The functions in Figure 3b begin to diverge at a contrast of around 4%, and data from individual participants showed a similar pattern (see Supplementary Figure S1).

**Figure 3:**
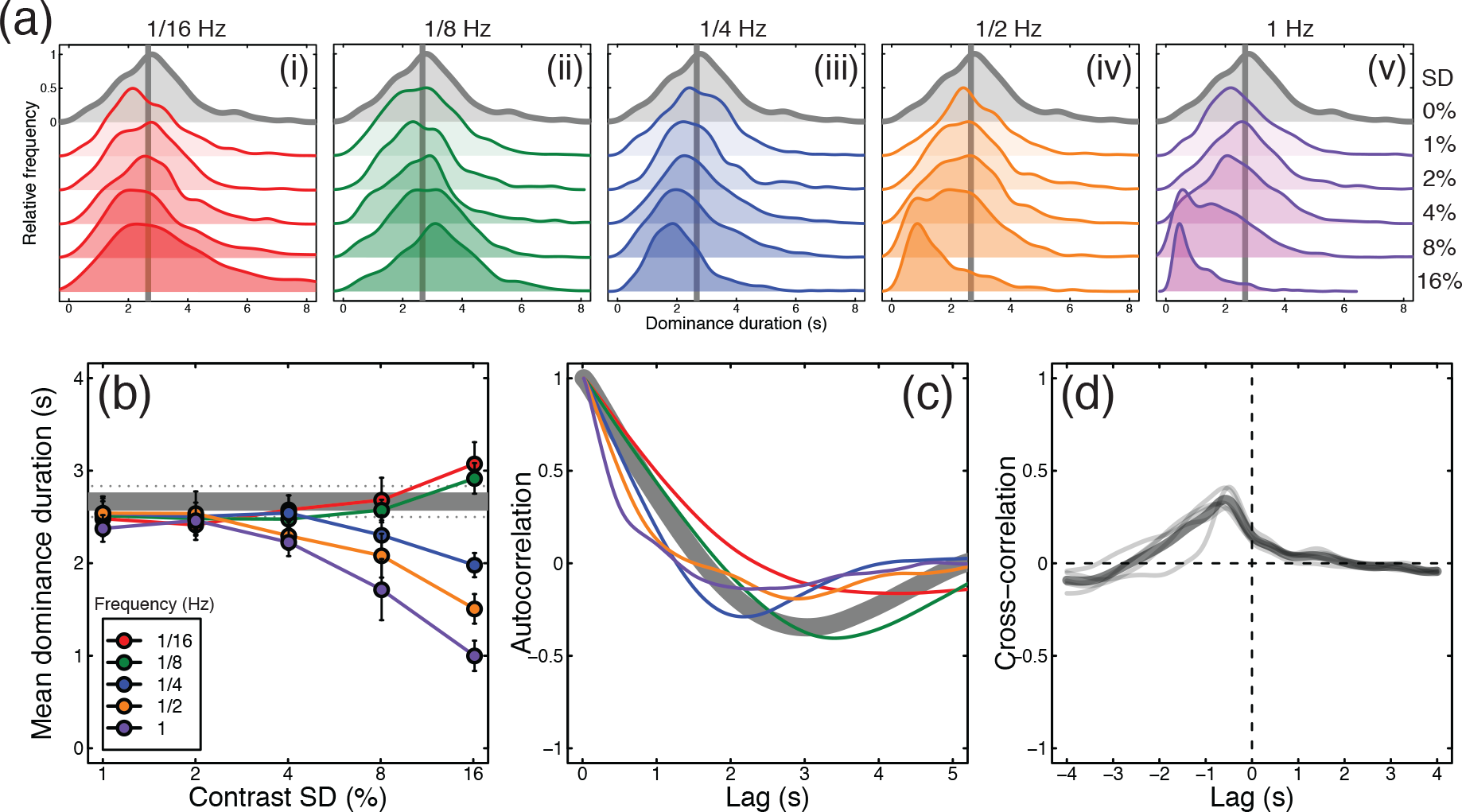
Traditional rivalry measures for all conditions, averaged (or pooled) across all participants (N=5). Panel (a) shows histograms of pooled dominance durations at five temporal frequencies (i-v) and a range of contrast levels (standard deviations of 0 − 16%, increasing down each plot). The grey histogram, duplicated in each plot, shows the baseline condition with no contrast modulation. For low temporal frequency, high contrast modulations, there were more very long dominance periods (the positive skew of the red histogram increases). For high temporal frequency, high contrast modulations there were more short dominance periods, and the histograms shifted left. Panel (b) shows mean dominance durations for all conditions, plotted as a function of modulation contrast. The grey horizontal line shows the baseline (no modulation) condition. Error bars (and dotted lines) show ±1SE across participants. Panel (c) shows autocorrelation functions averaged across participants for the baseline condition (grey curve) and the highest contrast modulation at each temporal frequency (curves, see panels a,b for colour legends). Panel (d) shows the cross correlation between the difference in noise modulations at the highest modulation contrast, averaged across all modulation frequencies. The thin grey lines denote individual participants and the thick black line is the average.

We also cross-correlated the noise time course (difference between left and right eye contrasts for the 16% contrast modulation conditions pooled across all temporal frequencies) with the participants’ responses (Figure 3d). This revealed a mean response lag of 583ms, somewhat faster than estimates from previous studies (Baker and Graf, 2009a). The mean cross-correlation coefficient at this time point was 0.35, indicating that a substantial proportion of the variance in participant percepts was predictable from the changes in stimulus contrast. Functions for individual participants are shown by the thin traces in Figure 3d, and are similar to the mean. Note that the auto-and cross-correlation functions shown here differ from the switch-triggered-average reverse correlation measure reported by Lankheet (2006), and the serial correlation measures used by Lehky (1995), van Ee (2009) and others (where ‘lag’ on the x-axis refers to dominance epoch rather than time). These measures assess different aspects of rivalry data that are not the focus of the current work.

Next, we calculated the consistency of responses across pairs of presentations of identical noise streams. In the absence of any noise modulation, the mean consistency was slightly above the expected baseline of 0.5, having a value of 0.53 (horizontal grey lines in Figure 4). The most likely explanation for this is that slight eye dominances or biases towards one or other stimulus will increase the consistency across repetitions, however the effect is very small. For conditions where the stimulus contrast was modulated, a 5 × 5 repeated measures ANOVA indicated that the response consistency depended on both temporal frequency (F(4,16)=9.90, *p*<0.001, *η*_*p*_^2^=0.71) and modulation contrast (F(4,16)=28.81, *p*<0.001, *η*_*p*_^2^=0.88), as well as the interaction between the two variables (F(16,64)=3.55, *p*<0.001, *η*_*p*_^2^=0.47). These effects are shown in Figure 4, which plots the same data twice as a function of either modulation contrast (Fig 4a) or temporal frequency (Fig 4b). The general trends are that consistency increases with contrast, and at each contrast is strongest for the 1/8Hz temporal frequency (shown in green). The maximum consistency was 0.72, for the 1/8Hz, 16% contrast condition, which is particularly noteworthy given that this temporal frequency had the weakest influence on dominance durations (see green points in Figure 3b). Consistency exceeded baseline for the 1/8Hz condition at around 4% modulation contrast (green diamonds in Figure 4). These main findings were also clear in the data of individual participants, shown in Supplementary Figure S1.

**Figure 4:**
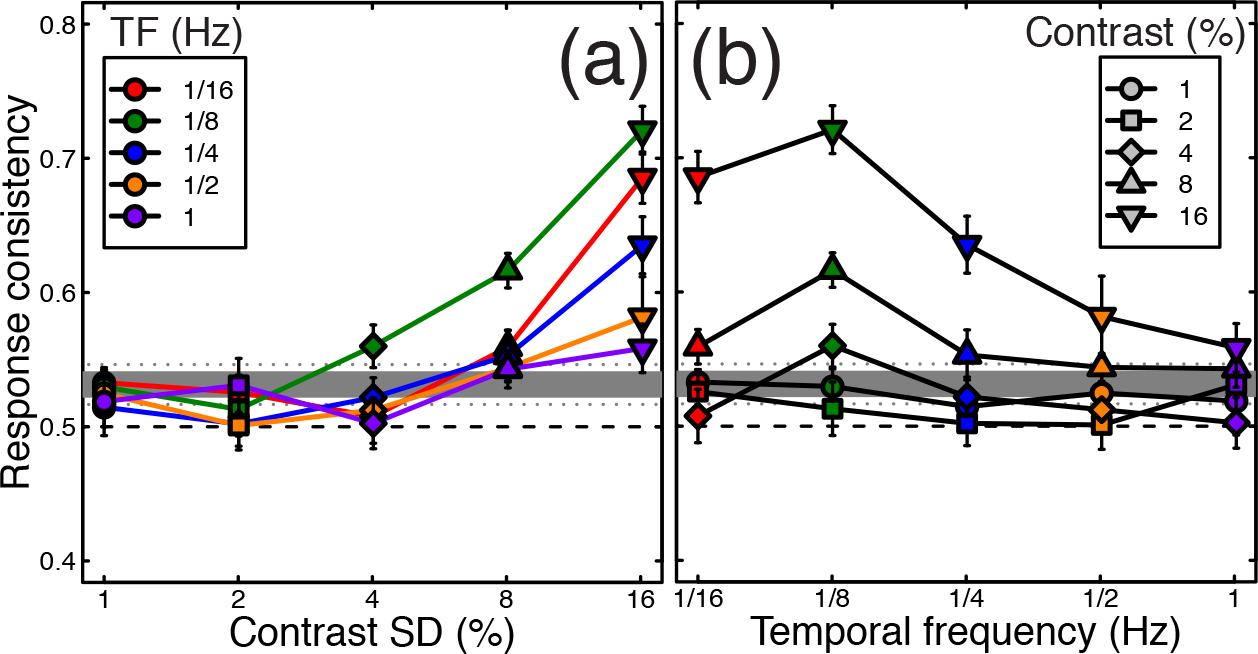
Response consistency across two passes through the experiment. The same data are plotted in both panels, as a function of modulation contrast (a) or temporal frequency (b). In each panel, the thick grey line represents the baseline (no modulation) condition, colours represent different temporal frequencies, and symbol types represent different contrasts. All data points are averaged across participants, with error bars indicating ±1SE of the mean. The dashed horizontal line at y=0.5 indicates a theoretical baseline in the absence of any response bias or eye dominance effects.

### A computational model with pink internal noise describes the human results

We first investigated how the amplitude of internal noise, and its spectral slope (α), affected model behaviour. We selected a single stimulus condition (stimulus noise frequency of 1/8Hz and amplitude of 16%) and ran the model with a range of internal noise contrast levels (SD = 1 − 64%) at five different spectral slopes (α = 0 − 2). The results of our simulations on dominance duration and response consistency are shown in Figure 5a(i-v), with the equivalent human data plotted in green for comparison. For all spectral slopes, as internal noise contrast increased it more strongly affected rivalry alternations. This is shown by the change in dominance duration (Figure 5b; increases for steep slopes and decreases for shallow slopes), and response consistency (Figure 5c), which decreased as responses became increasingly dominated by internal noise.

**Figure 5:**
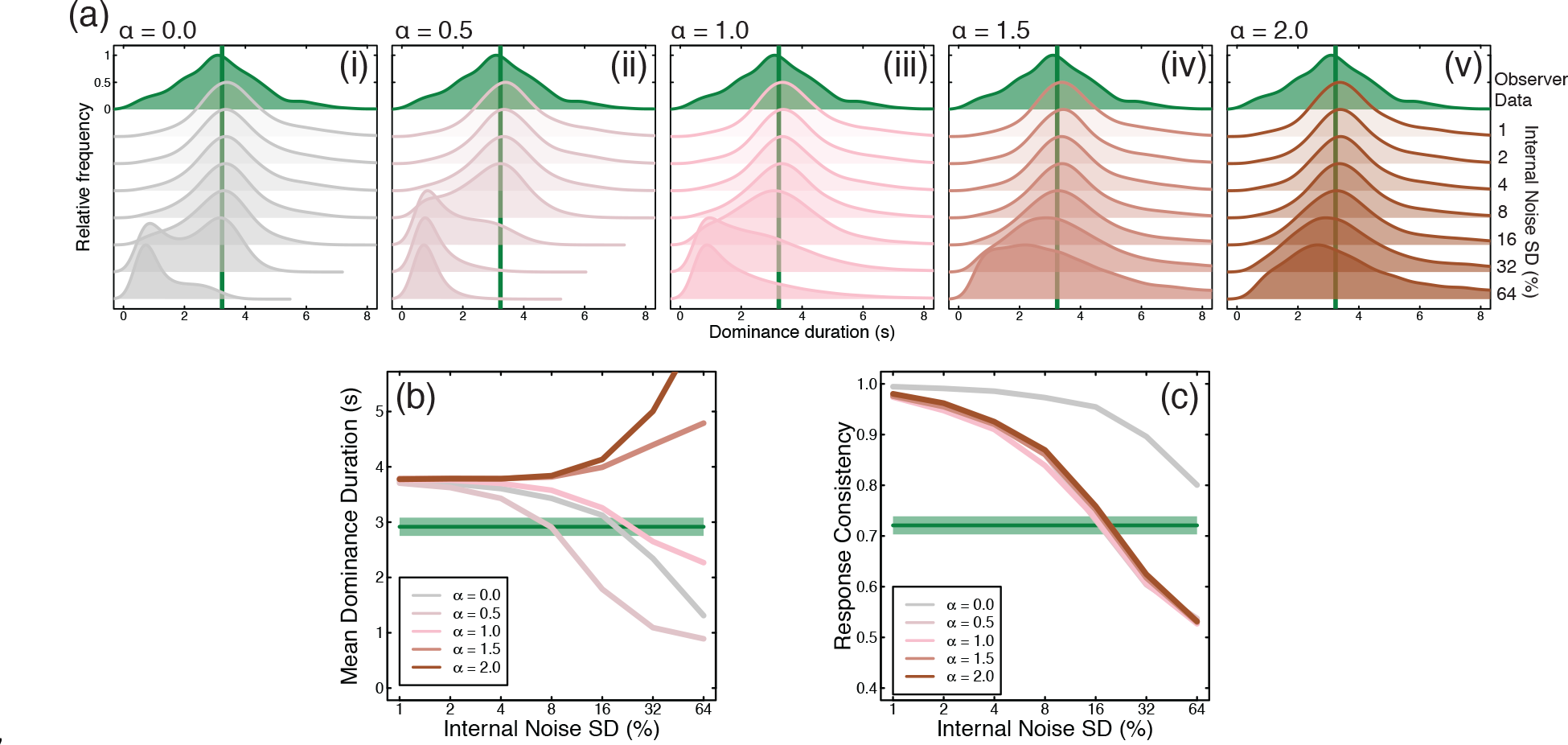
Summary of model behaviour for internal noise amplitude and spectral slope estimation. (a) The histograms of dominance durations for each spectral slope (α = 0.0 − 2.0) and contrast (SD = 1% − 64%) values. Within each subplot, the uppermost (green shaded) histogram shows the equivalent human data for a stimulus temporal frequency of 1/8Hz and a contrast modulation of SD = 16%. The solid vertical green line marks the average dominance duration for human observers. Histograms below show model dominance duration distributions for each internal noise contrast value. (b) Average dominance durations of the model for each spectral slope (coloured lines). The green line and shaded area mark human average dominance duration and ±1SE of the mean, respectively. Average dominance duration was affected by internal noise once its contrast reached 4%. Noise with steeper slopes (α = 1.5-2.0) increased mean dominance duration as a function of noise contrast, while noise with shallower slopes decreased mean dominance duration. (c) Response consistency decreased as a function of internal noise contrast for all spectral slopes. The green line and shaded area mark human observer average consistency and ±1SE of the mean, respectively. For all α>0, response consistency reached human levels at an internal noise contrast of 16%.

We can use the joint dominance durations and consistency scores to rule out several types of internal noise. White internal noise (α = 0) is not viable because there is no internal noise level for which both durations and consistency are close to human levels. Internal noise with steep amplitude slopes (α > 1) produces sensible consistency scores, but dominance durations become too long. This leaves slopes of α = 0.5 and α = 1, for which an internal noise contrast of around 16% gives a good approximation to the human data. We performed full simulations for all noise spectral slopes with this contrast. A slope of α = 1 was the best predictor of the human data, so these simulations are discussed in the main text, with simulations of other spectral slopes presented in Supplementary Figures S2 and S3.

The histograms of dominance durations, mean dominance duration and response consistency of the model simulations for all 26 stimulus conditions are shown in Figure 6. The model replicated the pattern of human data shown in Figures 3 & 4 remarkably well. The histograms of dominance durations of the model (Figure 6a i-v) show similar trends to those of human observers (Figure 3a). Slow modulation frequencies (1/16Hz and 1/8Hz) increased positive skew at high modulation contrasts (Figure 6a i-ii), while higher modulation frequencies shifted the peak of the dominance duration histograms leftwards as modulation contrast increased. The shifts in the histograms are reflected in the mean dominance durations of the model (Figure 6b), just as with human observers. Similarly, response consistency (Figure 6c, d) increased when stimulus noise contrast reached 4% and was highest for each contrast at a temporal frequency of 1/8Hz. Whereas human response consistency was quite bandpass (peaking at 1/8Hz and dropping quickly for faster frequencies), the model exhibited slightly broader tuning at high stimulus noise contrast. This may be due to the other parameters of the model that were fixed prior to our simulations, or it could imply additional physiological constraints such as bandpass temporal filters on the input, or variable response lag.

**Figure 6:**
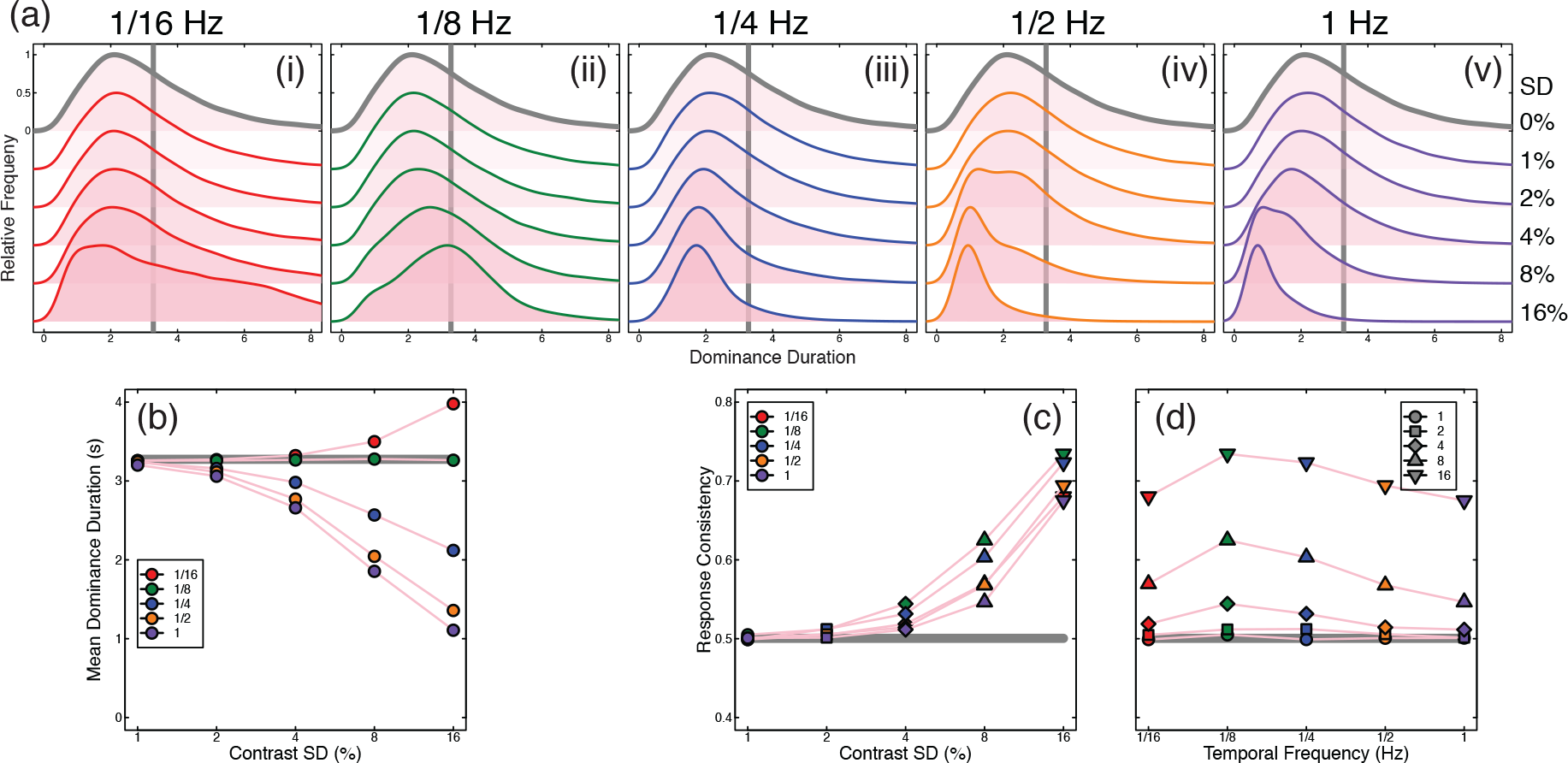
Summary of modelling results. (a) Histograms of dominance durations of the model with pink (α = 1) internal noise of 16% contrast for each stimulus temporal frequency (i-v) and contrast SD. The solid line colour serves as a legend for the stimulus noise temporal frequency (red = 1/16Hz, green = 1/8Hz, blue = 1/4Hz, yellow = 1/2Hz, purple = 1Hz). The histogram marked in grey represents baseline dominance durations with no contrast modulation. (b) The mean dominance durations of the histograms in (a). Marker colour represents the modulation temporal frequency, while the x-axis gives the modulation contrast. The grey line marks the baseline dominance duration of the model (3.18s), slightly slower than that of the human data. (c-d) Model response consistency plotted in the same manner as Figure 4. In (c), marker colour indicates the modulation temporal frequency while the x-axis indicates the modulation contrast. For all stimulus frequencies, response consistency increased according to modulation contrast, and was greatest when the stimulus temporal frequency was 1/8Hz. (d) Identical data but plotted with modulation temporal frequency on the x-axis. The grey line (c,d) marks response consistency at baseline with no external noise fed to the model (0.49).

### The model predicts consistency with antiphase noise sequences

We next explored whether the model could predict performance in novel conditions. Inspired by Kim et al. (2006), we designed a further condition in which the noise modulations were in antiphase across the eyes (i.e. a contrast increase in one eye matched with an equal contrast decrease in the other). We chose a temporal frequency of 1/8Hz, and tested four of our original participants. With no free parameters, the model described above made a clear quantitative prediction about performance in this condition (see Figure 7a), namely that response consistency should be reliably increased for the antiphase noise (brown squares in Figure 7a), compared to the equivalent condition from the main experiment with two independent streams of external noise (green circles in Figure 7a). This prediction was borne out empirically, as shown in Figure 7b. We note that dominance duration histograms from our human participants (and therefore mean dominance durations) remained relatively unaffected by this manipulation (see Figure 7c), consistent with performance with independent noise streams (Figure 3aii).

**Figure 7:**
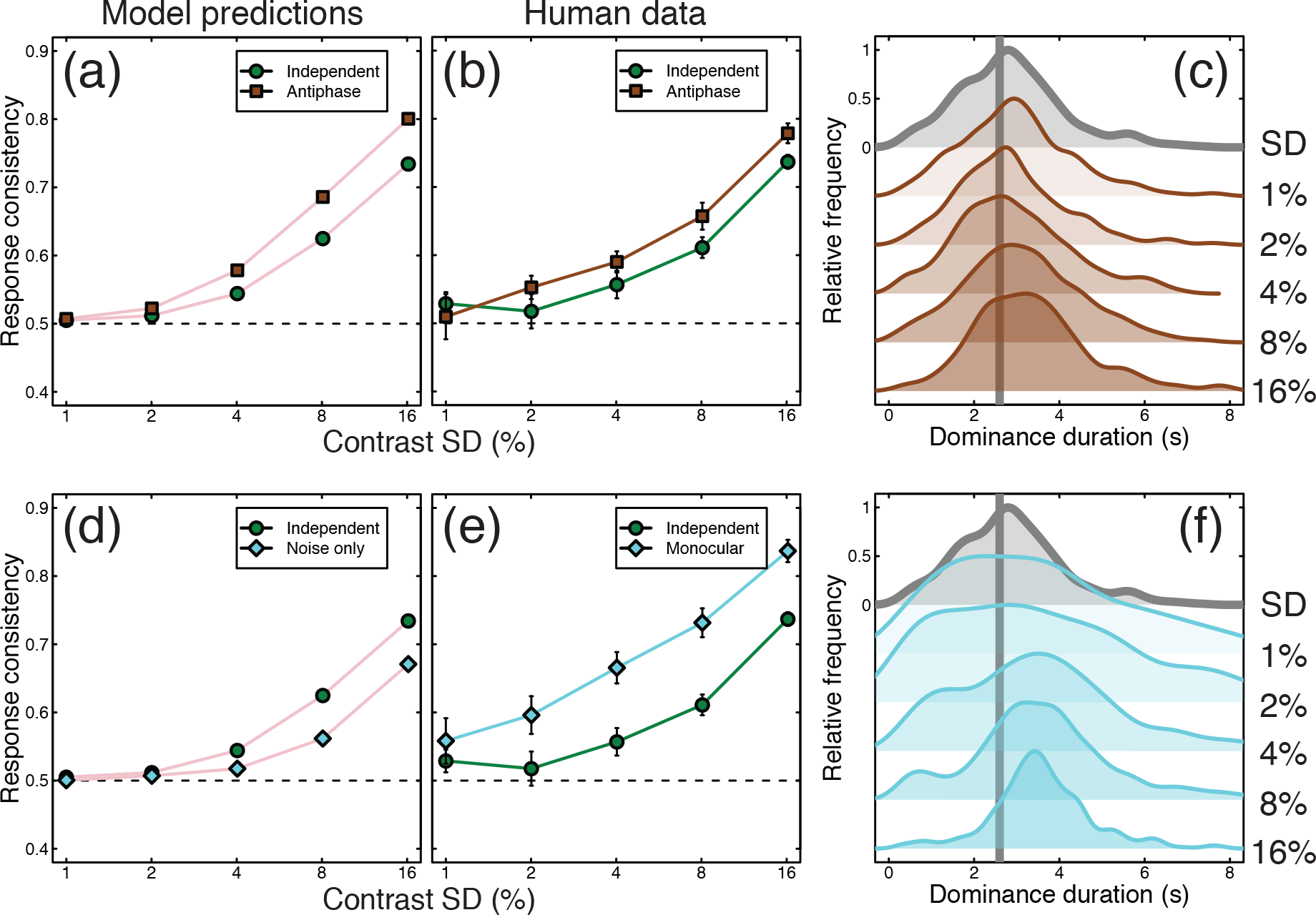
Summary of further conditions testing antiphase modulation and monocular rivalry. Panel (a) shows response consistency predictions of the model for independent (green circles) and antiphase (brown squares) external noise (modulation temporal frequency = 1/8hz). Panel (b) shows the human response consistency for the same conditions as (a). Panel (c) shows histograms of human dominance durations in the same format as Figure 3a, with the unmodulated rivalry condition shown at the top in grey. Panel (d) shows the response consistency of the model when the oscillatory mechanism is removed and modulations are driven by internal and external noise only (cyan diamonds) versus the response consistency for the main model (green cirxles). Panel (e) shows human response consistency to the ‘monocular rivalry’ condition (cyan diamonds) compared with that of the main experiment (green circles). Panel (f) shows human dominance duration histograms for the ‘monocular rivalry’ condition. Error bars and dotted lines show ±1SE across participants (N=4; for the conditions from the main experiment, we omitted results from the participant who did not complete the additional conditions when constructing this figure).

We also tested a condition in which we presented both stimuli to both eyes as a plaid, and modulated the contrast of the components. Just as in the main experiment, we asked participants to report which component appeared higher in contrast at each moment in time, though there was no binocular rivalry. This ‘monocular rivalry’ condition also produced greater consistency scores than the equivalent condition from the main experiment (see Figure 7e), and demonstrates that the technique can be used to dynamically monitor perception even in the absence of interocular competition. The distributions of dominance durations were rather broader for low contrast modulations, but narrowed at higher contrasts (see Figure 7f).

We reasoned that one way to model this condition might be to remove the rivalry mechanism from the model, leaving only the combination of internal and external noise to determine dominance at each moment. The predictions for this arrangement are shown by the cyan symbols in Figure 7d, and involve markedly *lower* consistency scores than both the model and empirical binocular rivalry conditions (green circles in Figures 7d,e), and also the monocular rivalry data itself (cyan diamonds in Figure 7e). Clearly then, monocular rivalry still involves some sort of alternation process (e.g. O’Shea et al. (2017), but see Georgeson (1984) for evidence to the contrary), but the increased empirical consistency scores in this condition suggest that the alternating mechanism is more strongly driven by the external noise modulations than during binocular rivalry.

## Discussion

Using a combination of psychophysical experiments and computational modelling, we infer that the source of internal noise relevant to perceptual alternations during binocular rivalry has an amplitude spectrum of 1/*f*, and a standard deviation of around 16%. Our method facilitates these inferences because it uses a double pass design, in which an external noise sequence is repeated twice, under the assumption that internal noise will be different on each pass. Although the double pass method has been used previously for briefly presented stimuli (Baker and Meese, 2012; Burgess and Colborne, 1988), this is the first time (to our knowledge) it has been used in a dynamic paradigm. We now discuss details of the rivalry model, relevance to other work on noise in binocular vision, and broader implications for our understanding of internal noise in the brain.

### Model variants and alternative models of rivalry

In the course of developing the model, we also considered several variants using same architecture that were either less successful or less plausible. One variant was a model in which a single source of internal noise was added to both channels. In this arrangement, the internal noise was less effective, because it increased or decreased the response in both channels by the same amount, and so did not materially influence the competition between channels. Another variant placed the internal noise sources outside of the gain control equation (i.e. added after eqn 1 rather than appearing on the numerator and denominator). Although moving internal noise later is consistent with the assumptions of a family of popular computational models of early binocular vision (Legge, 1984; Meese et al., 2006 see next section), this was less successful than our main model because internal noise levels sufficient to influence consistency had too large an effect on dominance durations. This rendered the dynamic properties of the model moot, with rivalry percepts being largely determined by the internal noise streams.

We also tested alternative values of the main parameters in the rivalry model. These altered model behaviour in the unmodulated baseline condition much as described in previous work (Wilson, 2007), but had relatively minimal effects on dominance durations and consistency scores with strongly noise-modulated stimuli, where rivalry alternations depend more on the interplay of internal and external noise than on adaptation and inhibition. We anticipate that other rivalry models with architectures related to that of Wilson (2007, 2003) could be modified in a similar way as described here to achieve comparable effects, but have not tested this assumption.

### Related work on rivalry

As mentioned above, Kim et al. (2006) modulated the contrast of rivalling stimuli periodically in antiphase at a range of temporal frequencies (building on earlier work by O’Shea and Crassini (1984) in which rivalling stimuli were entirely removed at different frequencies and phases). They implement three computational models to account for their results, each of which has random walk (i.e. brown) noise with a spectral slope of 1/*f*^*2*^, but report obtaining similar results with white noise for their experimental conditions. Furthermore, one of the models they implement is a version of the Wilson (2003) model considered here, but they report the best performance when the internal noise is added to the adaptation differential equation (see Methods), rather than the rivalling units (see also van Ee, 2009). In our simulations, we found similar effects on the dominance duration distributions for internal noise placed either in the main equation or adaptation equation (not shown here). However, placing internal noise in the adaptation differential equation resulted in response consistency that was not tuned to modulation frequency (i.e., flat). We suspect that Kim et al.’s paradigm did not afford sufficient constraints to distinguish between the two very different internal noise types or the locus of internal noise.

Other models that have incorporated a stochastic component include the model of Lehky (1988) which also used random walk (brown) noise, Kalarickal and Marshall (2000) who used additive uniformly distributed (effectively white) noise, and Stollenwerk and Bode (2003) who used temporally white noise that was correlated across space. A further model developed by Rubin and colleagues (Moreno-Bote et al., 2007; Shpiro et al., 2009) uses exponentially filtered white noise which progressively attenuates higher frequencies. However none of these studies report testing other types of internal noise, nor were their experimental conditions sufficient to offer meaningful constraints on the internal noise properties. As far as we are aware, this is the first study that has modelled internal noise of different amplitudes and spectral properties and compared the predictions to empirical results.

Baker & Graf (2009a) explored binocular rivalry using broadband pink noise stimuli that also varied dynamically in time. By testing factorial combinations of temporal amplitude spectra across the two eyes, they showed that stimuli with 1/*f* temporal amplitude spectra tended to dominate over stimuli with different spectral slopes (the same was also true of static stimuli with a 1/*f* spatial amplitude spectrum). Whilst these results do not directly imply anything about the properties of internal noise, they are consistent with the idea that the visual system is optimised for stimuli encountered in the natural world, which are typically 1/*f* in both space and time (e.g. Dong and Atick, 1995; Field, 1987; Geisler, 2008; Hansen and Essock, 2005; Simoncelli and Olshausen, 2001). Our findings here imply that as well as having a preference for external stimuli with naturalistic properties, the internal structure of the visual system might itself have evolved to emulate these temporal constraints (Field, 1987; Haun and Peli, 2013; Schwartz and Simoncelli, 2001; Schweinhart et al., 2017).

### Internal noise in binocular vision and throughout the brain

Early models of binocular signal combination attributed the improvement √2 in contrast sensitivity for fusible stimuli viewed binocularly vs monocularly to the pooling of independent monocular noise sources (Campbell and Green, 1965). However this model assumes that during monocular presentation, the noise in the unstimulated eye can be ignored, which is unlikely in the absence of experimental confounds (Legge, 1984). Contemporary binocular models of contrast detection and discrimination assume noise that is late and additive, occurring at a point beyond binocular signal combination (Meese et al., 2006). It is generally assumed that this late source of noise is the combination of multiple noise generators at successive stages of processing, though relatively little is known about their precise characteristics. However a small number of studies have investigated this issue, as we now summarise.

Pardhan & Rose (1999) added binocular external noise during a monocular or binocular detection task and found that binocular summation decreased at high levels of external noise, and that equivalent input noise (the minimum external noise level required to influence thresholds) was higher for monocular than binocular targets. One interpretation of these results is that the effective internal noise is greater for monocularly presented stimuli (see also Anderson and Movshon, 1989). However, the type of external noise that they used was broadband white pixel noise, which can also cause substantial gain control suppression (see Baker and Meese, 2012), potentially confounding the effects of increased variance. These results are therefore relatively inconclusive regarding sources of internal noise in binocular vision.

Recently, Ding & Levi (2016) have demonstrated that the inclusion of early (monocular) multiplicative noise in gain control models can account for some subtle features of binocular contrast discrimination performance. It has also been suggested that monocular noise might be increased in the affected eye of individuals with amblyopia (Baker et al., 2008). Finally, we have recently shown (Vilidaite et al., 2018) using a contrast discrimination paradigm that EEG and MEG data are consistent with both an early (~100ms post stimulus onset) noise source in low level visual areas, and a later noise source in more frontal and parietal brain areas, both of which affect perceptual decisions. All of these results are therefore consistent with an early monocular source of internal noise, as included in our model, but do not preclude the addition of later sources of noise which we do not consider here.

Regarding noise more generally, surprisingly few studies have addressed the spectral and distribution properties of internal noise using psychophysical methods. The default assumption is typically that internal noise is Gaussian (owing to Central Limit Theorem) and white. However, Neri (2013) concluded that internal noise had a Laplacian distribution, and other psychophysical work has assumed Poisson processes for internal noise (May and Solomon, 2015), based on single cell recordings (Goris et al., 2014). Noise with a pink amplitude spectrum typically retains a Gaussian distribution, though in principle non-Gaussian distributions (such as Laplacian or Poisson distributions) could also be altered to have a pink spectrum. Although we are unaware of any other psychophysical studies attempting to estimate the spectral characteristics of internal noise, we note that measurements of spontaneous neural activity using ECoG and fMRI also have fractal properties, and a slope of approximately 1/*f* in visual areas (He et al., 2010).

## Conclusions

Using a novel dynamic double pass paradigm with binocular rivalry, we measured how alternation rates and response consistency were affected by different types and amounts of external noise. The results were consistent with a computational model of rivalry in which internal noise was independent in each monocular channel. We conclude that internal noise relevant to rivalry has an amplitude spectrum of 1/*f*, and a standard deviation of around 16%. We anticipate that future studies might use temporally sensitive neuroimaging techniques such as EEG and MEG to further investigate these sources of internal noise.

## Methods

### Participants

The main experiment was completed by five psychophysically experienced observers (2 male), who provided written informed consent. Two were the authors, the remainder were unaware of the aims or design of the study. A control experiment was completed by four of the same observers. All observers had no known abnormalities of binocular vision, and wore their standard optical correction if required. Procedures were approved by the Ethics Committee of the Department of Psychology at the University of York.

### Apparatus and stimuli

Stimuli were sinusoidal grating patches with a spatial frequency of 1c/deg, subtending two degrees of visual angle, and ramped in contrast by a cosine function over a further ¼ degree. The gratings shown to the left and right eyes had orthogonal orientations (±45 degrees) which were assigned randomly on each trial (see Figure 1a for examples). The mean Michelson contrast of the gratings was 50%, but this was modulated by dynamic noise streams of various centre frequencies (1/16 Hz to 1Hz) and standard deviations (1% to 16% Michelson contrast). The noise streams were constructed by bandpass filtering white noise at the required frequency using a one octave bandpass filter (see Figure 1b). In the main experiment, the noise streams used to modulate the contrast of each eye were independent.

Stimuli were displayed on a ViewPixx 3D display (VPixx Ltd., Canada), driven by an Apple Macintosh computer. The monitor operated with 16 bits of greyscale luminance resolution (M16 mode) and was gamma corrected using a Minolta LS110 photometer. Independent stimulation of the left and right eyes was achieved using stereo shutter glasses (NVidia 3D Vision), synchronised with the monitor refresh rate of 120Hz via an infra-red signal. To promote good vergence and binocular alignment, each stimulus was surrounded by a static high contrast greyscale Voronoi texture (squares of 14 × 14 degrees, with a 7 degree diameter disc in the centre set to mean luminance) that was identical in both eyes (see Figure 1a). A different texture was presented on each trial, selected at random from a set of 99 pregenerated textures.

### Procedure

Participants sat in a darkened room and viewed the display from a distance of 57cm. Stimuli were presented for 60 seconds per trial, with condition order determined at random. Participants were instructed to indicate using a two-button mouse which of the two grating stimuli they perceived at each moment in time by holding down one or other button. If both stimuli were perceived, they were instructed to choose the stimulus that was most visible (i.e. that took up the largest part of the image), or to hold down both buttons if they were equally salient. At the end of each trial, there was a minimum blank interval of three seconds, with the following trial initiated by the participant.

Each of the 26 conditions (5 contrasts * 5 temporal frequencies + 1 baseline) was repeated 5 times by each observer using unique noise sequences in each repetition, and then a further 5 times using the same noise sequences as in the first pass. This resulted in 260 trials (4.3 hours of rivalry data) per participant, which were completed across multiple sessions (each typically lasting 20-30 minutes) over several days. Raw data are available online at: http://dx.doi.org/10.6084/m9.figshare.7262201

### Modelling

There are multiple models that have been successful at capturing the oscillatory behaviour of dominant percepts in binocular rivalry (Kim et al., 2006; Laing and Chow, 2002; Lehky, 1988; Wilson, 2007, 2003). While they vary in complexity, all include two key characteristics: inhibition between units responding to the left and right monocular stimuli, and self-adaptation. These guarantee that only one unit will be active at a given moment, and that over time, the active unit will decrease its firing rate sufficiently to allow the suppressed unit to be released from inhibition. Apart from a few exceptions (Brascamp et al., 2006; Kalarickal and Marshall, 2000; Kim et al., 2006; Lehky, 1988; Moreno-Bote et al., 2007; Shpiro et al., 2009; Stollenwerk and Bode, 2003), most computational investigations of binocular rivalry have focused on deterministic implementations of their models to investigate how suppression and self-adaptation contribute to oscillations in perceptual dominance. It is, however, fairly straightforward to adapt these models of rivalry to include an additive noise term and directly probe the properties (i.e., amplitude and spectral qualities) of internal noise. Here, we probe the properties of internal noise with the minimum rivalry model of Wilson (2007, 2003).

The minimum rivalry model defines the response of a single unit by two differential equations (equation 1 and equation 2), which incorporate stimulus excitation (*L*/*R*), self-excitation (*ε* = 0.2), competitive inhibition (*ω* = 3.5), self-adaptation (*H*), and here, an additive internal noise term (N). For the unit responding to stimuli presented to the left eye (E_L_), the response term is

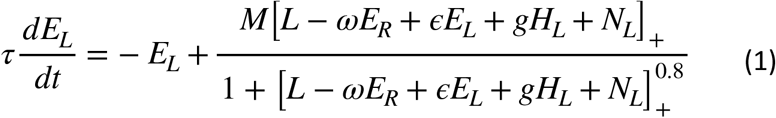

and self-adaptation is

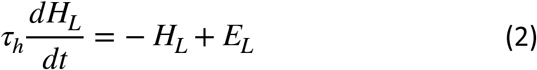

which is identical for activity in the right eye (ER), but with the subscripts switched. The constants *M* and *g* serve to scale the response gain and adaptation strength and were set to values of 1.0 and 3.0, respectively. The excitatory (τ) and hyperpolarizing (τ_*h*_) time constants of equation 1 and equation 2 were set to 15ms and 4000ms respectively. All model parameters were fixed in our simulations. Internal noise was additive and independently generated for each eye. As previous studies have already investigated the locus of internal noise with this particular model (Kim et al., 2006), we chose here to only conduct model simulations with noise added to the unit response equation (equation 1). Note that as the stimulus input to the model is identical to that of the psychophysical experiment (see Figure 2a), we use a contrast gain control variant of the Minimum rivalry model (Wilson, 2007) to account for any differences in contrast between eyes. This also means that the noise term is added to both the numerator and denominator of equation 1.

We probed the spectral characteristics of internal noise by injecting the model with broadband noise patterns (1/*f*^α^) generated at one of five different spectral slopes [^*^ We also conducted simulations with bandpass filtered internal noise streams with the same frequencies as that of the stimulus sequences, in addition to the broadband internal noise simulations. Response consistency was high for all stimulus conditions, which suggests that this type of internal noise is incapable of modulating model responses beyond that of the external noise sequences. As these results do not offer any additional insight to the characteristics of internal noise, we do not show them here.], where α = [0, 0.5, 1.0, 1.5, 2.0] (see Figure 2b). Noise patterns were generated in the Fourier domain by first creating a flat (α = 0) amplitude spectrum and then multiplying the amplitude coefficient at each frequency by *f*^−α^. The phase of each frequency component was assigned a random value between -π and π. Two different phase spectra were generated in order to create two independent noise streams (N_L_ and N_R_) with the same amplitude spectrum. These were rendered in the temporal domain by taking the inverse Fourier transform and adding them to the left and right units separately.

Perceptual switches were implemented as a winner-take-all rule: the dominance of a percept was defined by the magnitude of E_L/R_ at any given moment in time (if E_L_ > E_R_, E_L_ is dominant; see Figure 2c) Finally, all model simulations were conducted in MATLAB (version R2017a) using ODE45 to solve the 4 differential equations that define the response of each unit and their self-adaptation over 60 seconds (i.e. the duration of a trial in the psychophysical experiment). We simulated binocular rivalry twice - with different internal noise samples but the same external noise sequences – for each stimulus noise condition in order to calculate the response consistency of the model. This was repeated 1000 times, and the model outputs (dominance duration and response consistency) were averaged across repetitions.

## Acknowledgements

Supported in part by the Wellcome Trust (ref: 105624) through the Centre for Chronic Diseases and Disorders at the University of York. We thank Robert O’Shea and two anonymous reviewers for helpful comments on a previous version of the manuscript.

## Supplementary information

**Figure S1.**
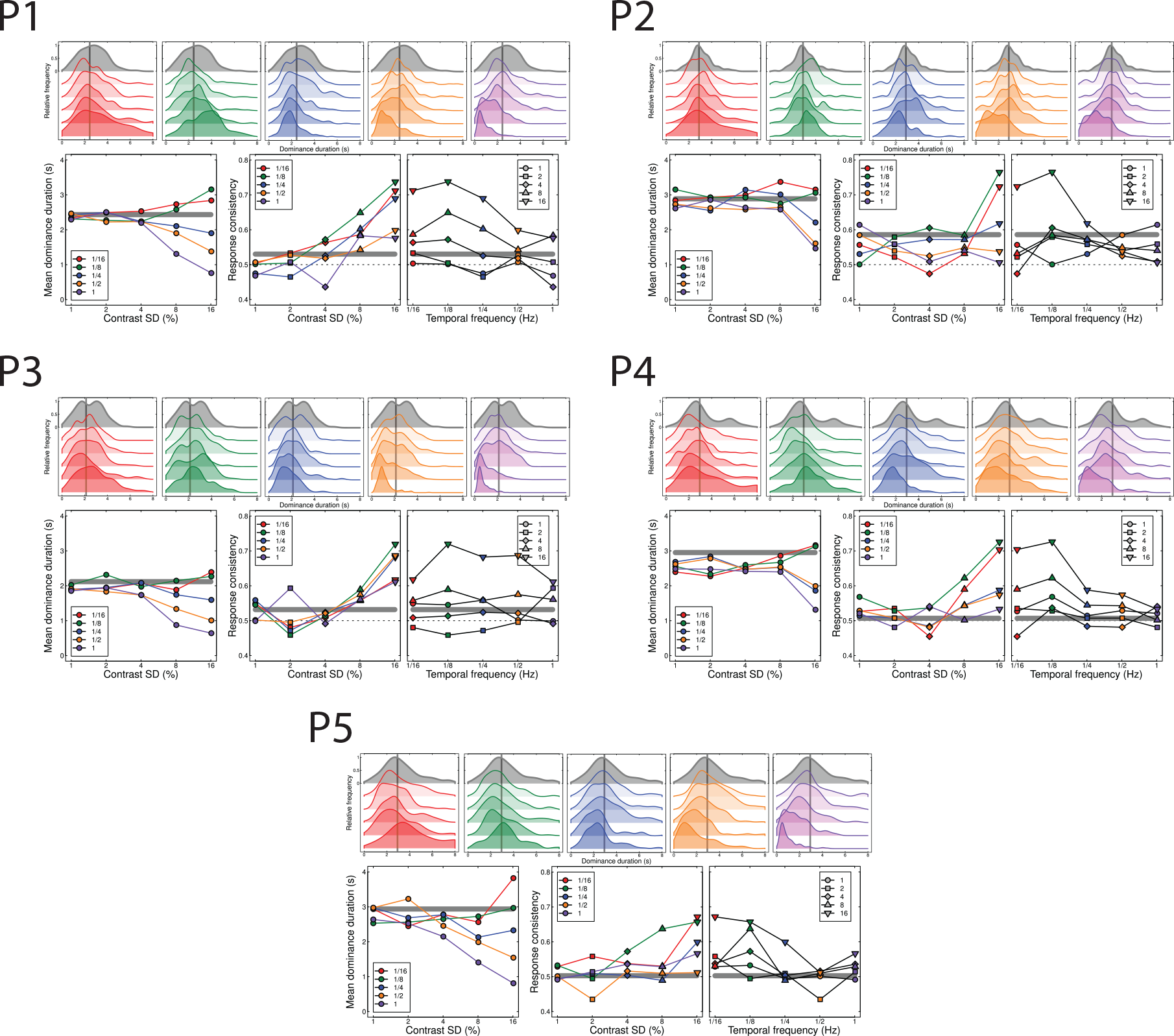
Data for individual participants (P1-5). See the captions to Figures 3 and 4 for formatting details

**Figure S2.**
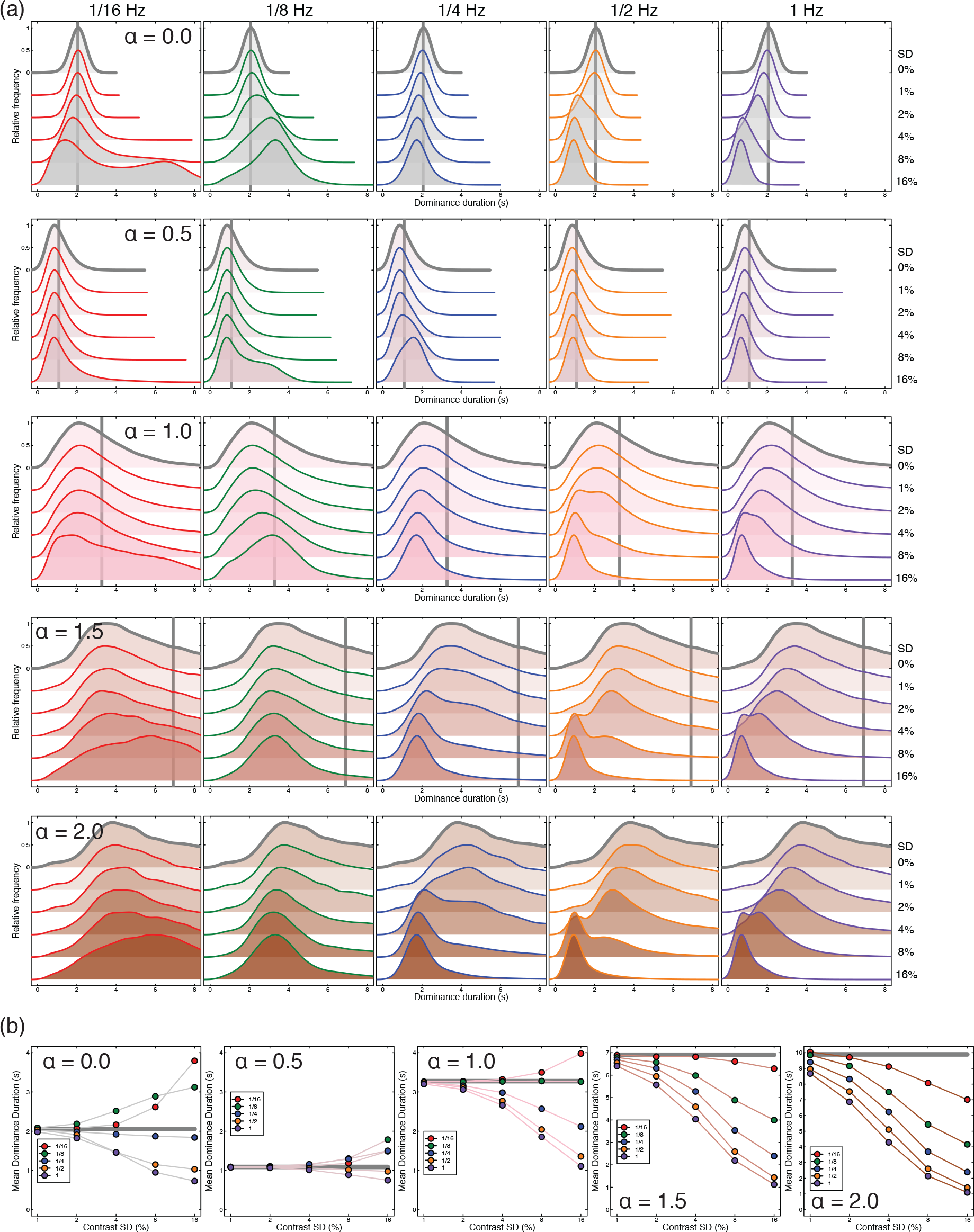
(a) Model dominance duration histograms for each of the five noise αs and stimulus condition as in Figure 6a. The solid line colour indicates the stimulus temporal frequency while the fill colour marks the noise α. The grey vertical line marks the mean dominance duration of the 0% modulation contrast condition. For very steep slopes (α = 2) the mean exceeds the x axis limit (~10s). (b) The average dominance duration for each model noise α as in Figure 6b. Note the different scale for the y axis with internal noise αs of 1.5 and 2.0.

**Figure S3.**
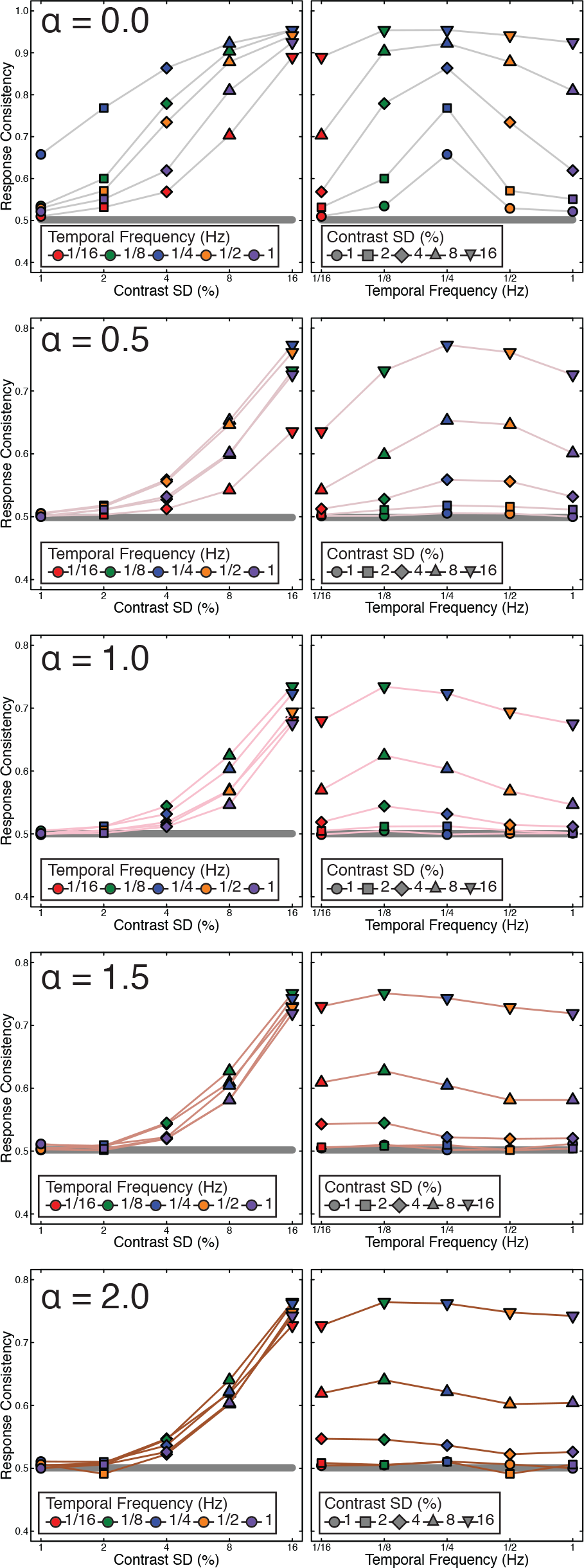
Response consistency for all five internal noise α values investigated here. The left column charts response consistency for each modulation contrast while the right column shows the same data replotted according to modulation frequency as in Figure 6c and 6d.

